# Partially methylated domains are hallmarks of a cell specific epigenome topology

**DOI:** 10.1101/249334

**Authors:** Abdulrahman Salhab, Karl Nordström, Kathrin Kattler, Peter Ebert, Fidel Ramirez, Laura Arrigoni, Fabian Müller, Cristina Cadenas, Jan G. Hengstler, Thomas Lengauer, Thomas Manke, DEEP Consortium, Jörn Walter

## Abstract

**Background:** Partially methylated domains, PMDs, are extended regions in the genome exhibiting a reduced average DNA-methylation level. PMDs cover gene-poor and transcriptionally inactive regions and tend to be heterochromatic. Here, we present a first comprehensive comparative analysis of PMDs across more than 190 WGBS methylomes of human and mouse cells providing a deep insight into structural and functional features associated with PMDs.

**Results:** PMDs are ubiquitous signatures covering up to 75% of the genome in human and mouse cells irrespective of their tissue or cell origin. Additionally, each cell type comes with a distinct set of specific PMDs, and genes expressed in such PMDs show a strong cell type effect. Demethylation strength varies in PMDs with a tendency towards a more pronounced effect in differentiating and replicating cells. The strongest demethylation is observed in highly proliferating and immortal cancer cell lines. A decrease of DNA-methylation within PMDs tends to be linked to an increase in heterochromatic histone marks and a decrease of gene expressions. Characteristic combinations of heterochromatic signatures in PMDs are linked to domains of early, middle and late DNA-replication.

**Conclusion:** PMDs are prominent signatures of long-range epigenomic organization. Integrative analysis identifies PMDs as important general, lineage- and cell-type specific topological features. PMD changes are hallmarks of cell differentiation. Demethylation of PMDs combined with increased heterochromatic marks is a feature linked to enhanced cell proliferation. In combination with broad histone marks PMDs demarcate distinct domains of late DNA-replication.

## Background

DNA-methylation is an epigenetic hallmark with an important role in gene and genome regulation. Changes in the genome-wide landscape of DNA-methylation are extensively studied in the context of small regulatory regions like CpG islands [1], CpG shores [2], proximal and distal regulatory regions [3]. With the first genome-wide bisulfite-based DNA methylation analyses, a new term, partially methylated domains (PMDs), was introduced by Lister et.al. [4], referring to long genomic regions in the range of hundreds of kilo-basepairs (kb) characterized by highly disordered methylation levels. They were initially discovered in the fibroblast cell line IMR90 but cannot be observed in human embryonic stem cells H1.

It has been shown later that PMDs are enriched with heterochromatic histone modifications such as H3K27me3 and that they are gene-poor and less active [5, 6] than other genomic regions. Several studies have since reported PMDs in various cell types: medulloblastoma [6], adipocyte tissue [7], SH-SY5Y neuronal cells [8] and human cancers [5, 9, 10, 11]. PMDs in cancer cells are linked to late replication and nuclear lamina associated regions [10]. The first non-cancer primary human tissue type with PMDs has been reported in placenta [12]. Recently we, as part of the DEEP consortium http://www.deutsches-epigenom-programm.de/, published the first primary human cells, CD4+ T cells, with PMDs [13]. We showed that progressive loss of DNA methylation correlates with T cell memory differentiation and happens predominantly in PMDs. Burger et al. [3] implemented an HMM-based detection method called MethylSeekR to define PMDs and separate them from fully methylated regions (FMRs) and short (regulatory) regions that come in two types; lowly methylated regions (LMRs, CpG poor regions) and unmethylated regions (UMRs, mostly CpG islands). LMRs and UMRs are relatively short (a few hundred to a few thousand basepairs) and correspond to distal and proximal regulatory elements, respectively [14]. Tools such as MethylSeekR are very useful for exploring the methylome landscape on a large scale and help to discriminate the large domains, from the small regulatory regions.

In collaboration with our colleagues in the international human epigenome consortium IHEC http://ihec-epigenomes.org/, we contributed to generating a large epigenome cohort for numerous primary cell types from human and mouse. WGBS data serve as an invaluable resource for studying PMDs in primary cells. PMDs represent a new aspect for studying the methylation landscape on a genome-wide level apart from the context of regulatory regions that have been studied extensively and pose the question whether DNA-methylation has an impact on the genome organization. At the same time, it has become quite clear that cells *in vitro* behave differently from primary cells, for instance regarding methylation levels. Thus it is important to compare the methylome of primary cells and cell lines in order to validate *in vitro* systems and afford an appropriate interpretation of the data.

Here, we investigate the genome-wide organization of PMDs across a comprehensive spectrum of available WGBS data generated by IHEC members; DEEP (http://www.deutsches-epigenomprogramm.de/), Blueprint (www.blueprint-epigenome.eu) and Roadmap (http://www.roadmapepigenomics.org/), together with other public data in order to gain insights into PMDs. In addition, we integrated WGBS data with other epigenetic data; ChIP-seq, RNA-seq, Hi-C and Repli-seq, in an attempt to describe the interaction between DNA methylation and chromatin formation in order to understand how they impact cell division, differentiation and the higher order chromatin structure. Moreover, we propose a new integrative approach to exploring and interpreting methylome topologies using WGBS data, an approach very much needed as the amount of such data is growing rapidly.

## Results

### Partially methylated domains are cell type discriminators

We collected and surveyed 171 public human WGBS data sets of different primary cell types (hepatocytes, T cells, B cells, monocytes, macrophages, eosinophils, neutrophils, dentritic cells, natural killer cells, endothelial cells and thymocytes) and tissues (liver, intestine, spleen, esophagus, stomach gastric, colon sigmoid, colon mucosa, heart and pancreas) for which we identified PMDs with MethylSeekR (see Additional file 1 for the complete list of samples). The lengths of PMDs vary broadly, ranging from 100kb up to 20Mb (Figure S1). PMDs cover a large portion of the genome (50 – 75%). The average and individual levels of PMD methylation vary between different cell types. While PMD positions in the genomes are highly conserved across cell types, in general, only roughly 26% of the genome is annotated as completely shared PMDs across all cell types (Figure 2B). Overall, PMDs are enriched for the broad heterochromatic marks H3K27me3 and H3K9me3 and depleted for the broad euchromatic mark H3K36me3. The latter is also reflected in the low appearance of annotated transcriptional units within PMDs and an overall low average transcription of genes located in PMDs (Figures 1 and S4).

**Figure 1.**
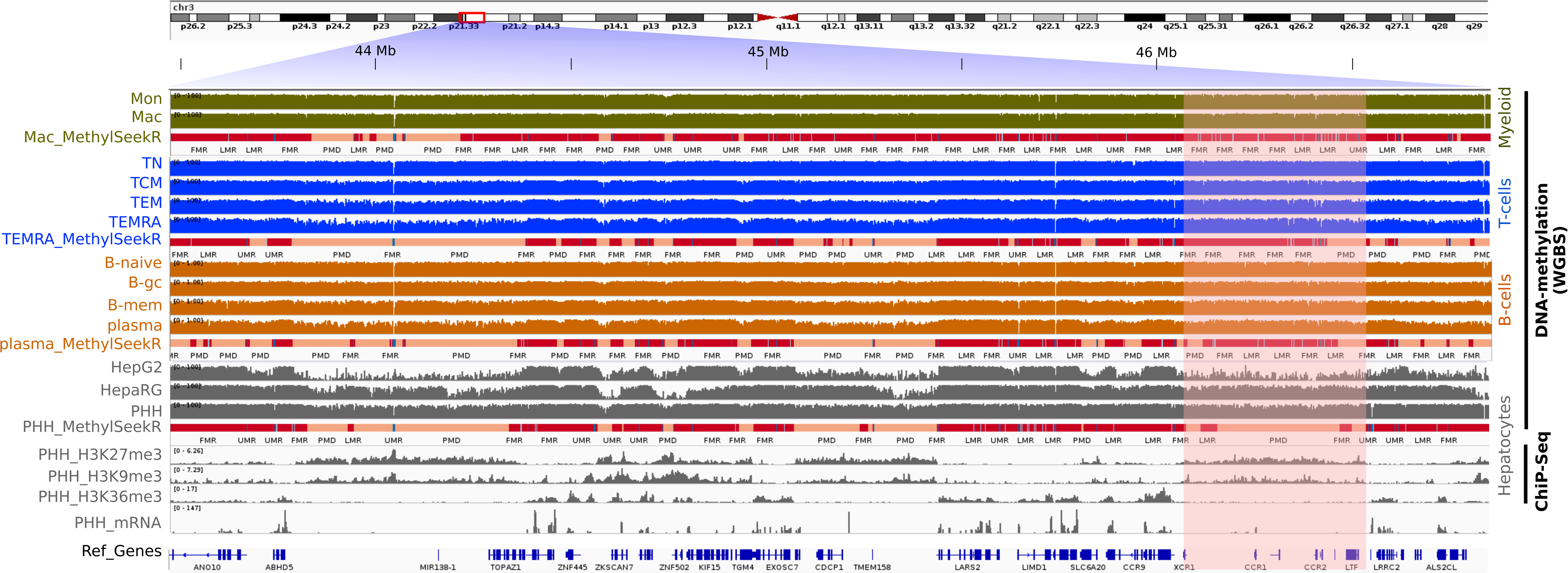
Genome wide DNA-methylation segmentation across different cell types. IGV snapshot showing DNA-methylation profiles of different cell types (from top to bottom: monocytes, macrophages, naive/central/effector/terminal memory T-cells, naive/germinal center/class switched memory B-cells, plasma, HepG2, HepaRG and hepatocytes) with the corresponding MethylSeekR segments: fully methylated regions (red); FMRs, partially methylated domains (light pink); PMDs, low methylated regions (light blue); LMRs and unmethylated regions (blue); UMRs (only one track per cell type shown for simplicity). Below the block of methylation data three broad histone marks and RNA-seq profiles of hepatocytes are displayed. PMDs can be seen as long regions with highly disordered methylation levels and tend to be largely overlapping between the different cell types. However there are cell type-specific PMDs (the highlighted region shows a hepatocyte-specific PMD). PMDs are gene-poor and transcriptionally inactive regions and have heterochromatic signature (H3K27me3 and H3K9me3). In contrast, FMRs are transcriptionally active and rich gene regions with enrichment of active histone mark H3K36me3.

To gain a deeper insight into the cell-specific and genome-wide distribution of PMD methylation profiles, we generated and applied a modified ChromHMM [15] approach, “ChromH3M”, as an abbreviation for ChromH**MM m**eta-segmentation (details in Methods and Figure S2). In brief, we bin the genome into 1kb tiled windows, labeled as 1 or 0 according to the presence/absence of PMDs for each sample. This binarized signal is then processed with ChromHMM to generate a fifteen-state model. The emission probabilities are displayed after hierarchical clustering. This approach generates PMD clusters discriminating cell type origin and/or cell-related subgroups (Figure 2A). Only five out of 171 samples did not cluster together with samples of similar origin. This approach is surprisingly stable even across cells which differ strongly in their overall methylation level (shown as box-plots in Figure 2A). We also used shorter LMR and UMR regions for such a ChromH3M meta-segmentation and roughly obtained the main subgroups in hierarchical clusters using 10,000 bootstraps and an “au” threshold of 97 (see Figure S3 and Methods for details). We conclude that PMDs are strong cell-type specific discriminators comparable with regulatory changes in short UMRs/LMRs.

**Figure 2.**
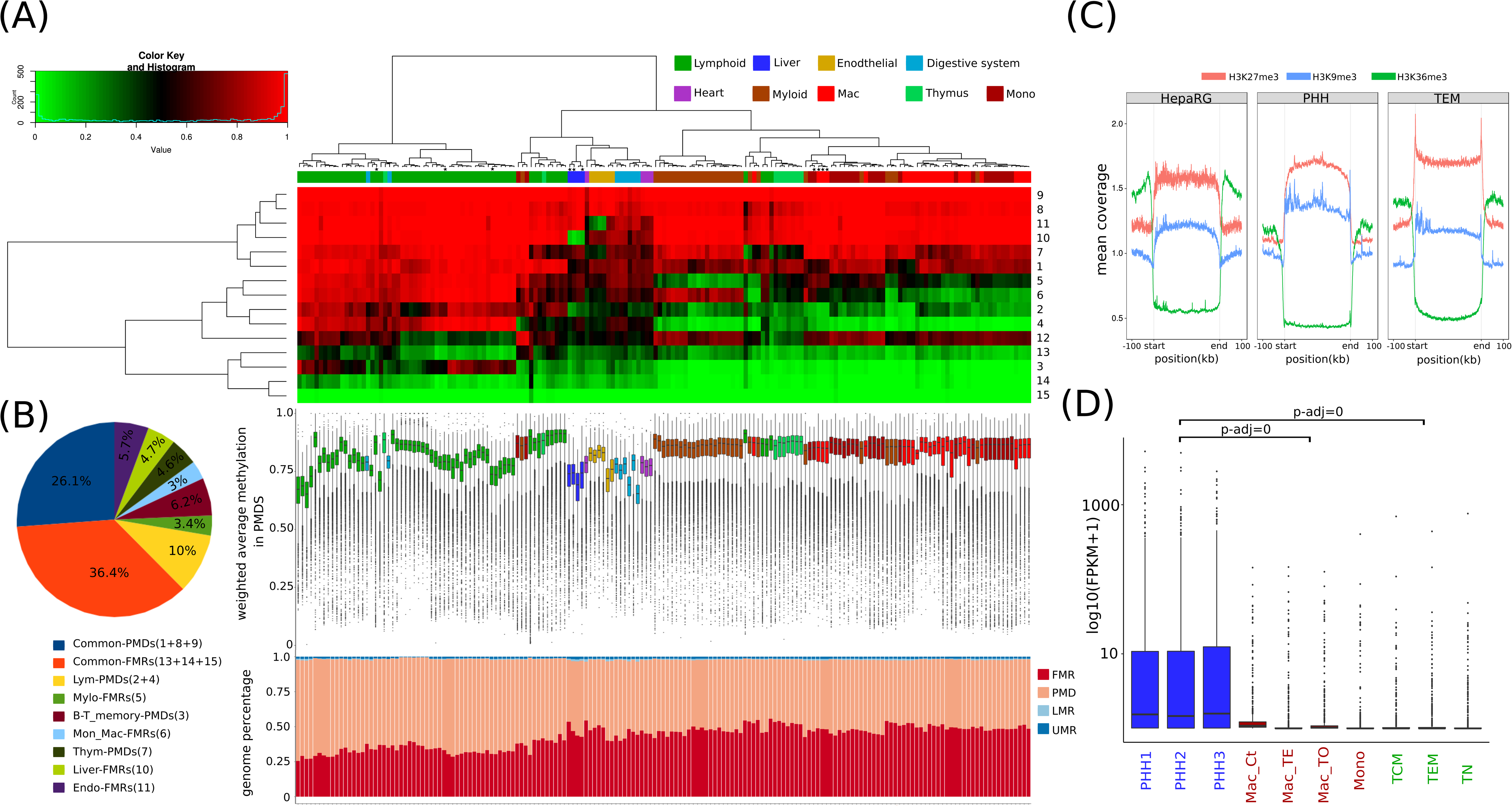
PMD signatures discriminate cell type and global transcriptional control. (A) Colored representation of the emission probabilities calculated by ChromH3M. Samples and states were hierarchically clustered forming six main groups; myeloid, lymphoid, endothelial, liver, digestive system and heart. Beneath each sample, the corresponding average methylation levels within PMDs is shown as whisker box plots and the percentage of MethylSeekR segments as stacked bar plots. Samples derived from the same cell type clustered together, although they differ in mean methylation level, suggesting that they have more similar PMD structure than the other cell types. PMDs comprise about 50 - 75% of the genome. (B) Graphical representation of the relative (percentage) contribution of each state in the 15 state ChromH3M model. 26% of the genome shares the same PMDs across all samples and roughly one third differ between them. (C) Genome-wide normalized histone mark signals within PMDs (including 100kb flanking regions). Note the enrichment of heterochromatic marks H3K27me3 and H3K9me3 across PMDs and a depletion of the transcription-coupled mark H3K36me3. (D) Log10-scaled FPKM values of state10 (Liver-FMRs) associated protein-coding genes. Genes are significantly more highly expressed in hepatocyte samples (PHH) than in macrophages, monocytes and T-cells, according to two-way ANOVA and Tukey HSD post-hoc test (details in Methods). Only samples marked with star in Figure 2A are used for simplicity and since they belong to one consortium.

For 171 available human methylomes of tissues and primary cells, ChromH3M generates a tree with six main branches separating myeloid from lymphoid cells, endothelial tissues, liver tissue, tissues of the digestive system and heart. Myeloid cells split into two subclusters; the granulocytes (neutrophils and eosinophils) and agranulocytes (monocytes, macrophages and dentritic cells). Members of both subclusters have similar average PMD methylation. B-cells and T-cells form a lymphocyte cluster which branches off into subgroups of memory T and B cells; i.e, central and effector memory T cells, germinal center and memory B-cells, respectively. This indicates that cell types not only display a distinct overall PMD topology but also acquire distinct PMD substructures upon proliferation and differentiation [13].

We furthermore observe that, in general, PMDs have extended heterochromatic signatures in both primary cells and permanent cell lines (Figure 2C, see also [4]). PMDs cover relatively gene-poor regions with mostly lowly/unexpressed genes (Figure S4).

The ChromH3M analysis reveals a couple of distinct features of cell-type specific PMDs (Figure 2A). For instance, state 10 and 11 comprise regions that only are FMRs in liver and endothelial cell types, respectively. State 4 discriminates myeloid FMRs from PMDs in other cell types. State 3 defines B and T cell specific PMDs (Figures 2A and 2B). The shared, i.e., common PMDs are defined by states 1, 8 and 9, while states 14 and 15 define common FMRs.

To explore the biological functions of genes present within cell-type specific FMRs/PMDs, we performed a functional annotation analysis with DAVID [16, 17] for genes in state 10 and state 3. For the former, liver-specific FMRs, the GO terms liver tissue expression, Rotor syndrom disease (lack of hepatocyte pigment deposits) and the KEGG pathway for drug metabolism through cytochrome P450 were obtained. These genes exhibit significantly higher expression in liver tissue/hepatocytes than in other cell types (Two-way ANOVA and Tukey HSD post-hoc test, p-adj=0) (Figure 2D). Furthermore, these FMRs are largely devoid of heterochromatic marks and enriched for the transcriptional elongation mark H3K36me3 across gene bodies (Figure S5, left panel). This is exemplified by two hepatocyte-specific gene loci CYP2B6 and FMO6P (Figure S6). The latter state, number 3, marks B- and T-cell-specific PMDs. Hence, these regions in B- and T-cells are enriched with the repressive mark H3K27me3 and, to a lower degree, with H3K36me3. Further, the functional analysis provides cell-type associated terms; cell differentiation, inflammatory response, adaptive immune response and specific surface antigen MHC class I, in addition to the KEGG pathway for the hematopoietic cell lineage. Interestingly, the expression levels of these genes are down-regulated in accordance with their PMD annotation. However, regarding only the methylation signal, there is a trend to split the B- and T-cells into naive versus memory cells. This discrimination can neither be confirmed by ChIP-seq nor by RNA-seq (see Figure S5, right panel). This could be due to the limitation in detecting the precise boundaries of shallow PMDs in naive cells.

In summary the ChromH3M results indicate a domain-wide transition of cell-type specific PMDs into FMRs and vice versa along with transcriptional regulation. The direction of this transition couples with specific changes in heterochromatic states.

A ChromH3M analysis on 24 WGBS mouse samples (Figure S7) shows a similar classification and distribution of PMD states, confirming that our findings hold for not only human but describe a feature apparently conserved among mammals. In mouse we identify cell-type/tissue specific PMDs for neuron, intestine, colon and mammary epithelial cells. Furthermore, the epithelial cells group into cells of the luminal and the basal compartment. We conclude that in human and mouse PMDs are excellent epigenome classifiers of cell-type specific topologies.

### Chromatin compaction increases with DNA-methylation erosion at PMDs in immortalized cells

Immortalized cell lines are widely used for studying cellular mechanisms including the influence of epigenetic control. However, it is known that cells in culture undergo drastic epigenetic alterations linked to passaging and cell replication numbers [18]. To investigate the epigenome wide changes occurring between primary cells and immortal cell lines, we compared the methylomes of primary cells and cell lines of the same origin. With this comparison we wanted to monitor the impact of cultivation and cancer-specific changes on PMD formation. We generated epigenome data for isolated primary hepatocytes (PHH) and two hepatic cancer cell lines; the hepatic progenitor cell line (HepaRG) and the liver hepatocellular carcinoma cell line (HepG2). We also include in our comparison results on publicly available liver cancer cells and noncancerous liver tissues.

First, we calculated the average methylation across the samples in 10Kb bins. We then performed k-means clustering forming six disjoint clusters which were subsequently annotated by the MethylSeekR segmentation (Figure 3B). Cluster 1 defines fully methylated bins across all samples while the other clusters show progressive loss of methylation in the order: liver tissue > PHH > liver cancer > HepaRG > HepG2. Interestingly, primary liver cancer was more similar to the noncancerous liver and PHH than to the cancer cell lines HepaRG and HepG2. Both cancer cell lines have lower methylation levels compared to the primary cells, as seen in clusters 4 and 5. This indicates a different epigenomic pattern in cultivated cancer cells in comparison to primary cancer cells. To gain deeper understanding of the features governing the development and changes in cell-type specific PMDs we focused on the analysis of liver PMDs that exhibit changes in the average methylation level in the cancer cell lines. We first extracted PMDs of PHH cells, which exhibit large overlap with PMDs of liver tissue and liver cancer tissue (data not shown). Such primary liver cell PMDs split into three subclasses, with respect to changes in DNA-methylation in HepaRG and HepG2 (Figure 3C). In the first subclass (class_I, red), PHH and HepaRG exhibit the same average degree of methylation (65%), but show a very low methylation state in HepG2. In the second class (class_II, green), HepaRG methylation levels are intermediate between PHH and HepG2, while in the third class (class_III, blue) both HepaRG and HepG2 show the same low average methylation as compared to the primary cells.

**Figure 3.**
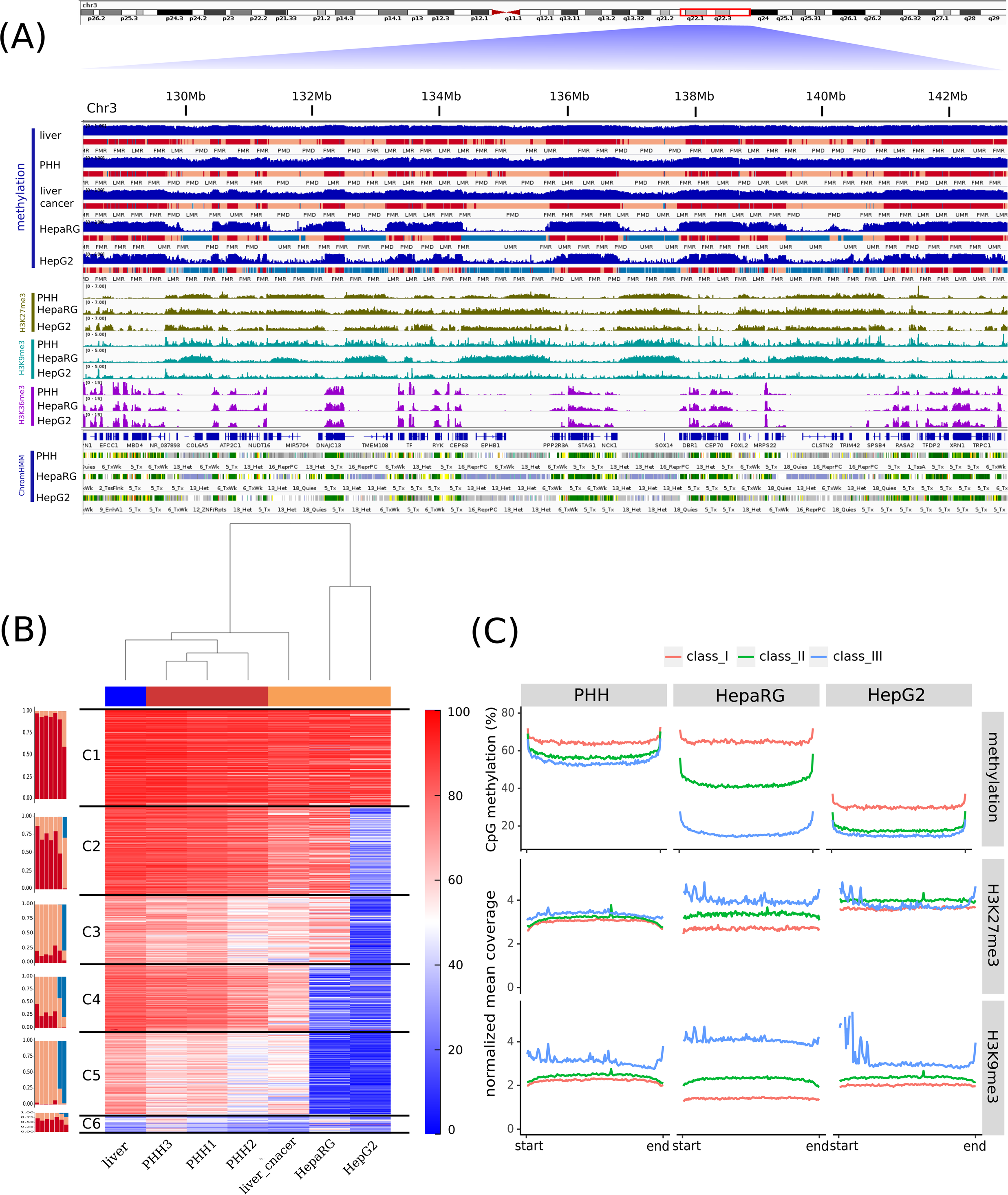
Heterochromatization accompanied by DNA-methylation erosion at PMDs in cancers. A snapshot of 14 Mb of chr3 showing the relevant epigenetic marks. Top: distinct raw DNA-methylation tracks and the MethylSeekR segmentation of liver tissue, isolated hepatocytes (PHH), liver cancer tissue, HepaRG and HepG2 cells lines, respectively. PMDs of primary cells, normal and cancerous tissues are extensively and selectively demethylated in cancer cell lines (largely converted into unmethylated regions). Middle: Histone marks H3K27me3, H3K9me3 and H3K36me3 in the same samples. Bottom: ChromHMM segmentation based on these six histone modifications and Input (details in Methods). (B) K-means clustering (k=6) based on the averaged methylation in 10 Kb bins. Cluster 1 represents the most (almost fully) methylated bins across all samples, while the other clusters are ordered according to the progressive erosion of methylation in PMDs. Bar plots (left) beside the heatmap show the percentage of the annotated bins as FMR, PMDs and UMR for each sample in each cluster. (C) Progressive DNA-demethylation in PMDs across cancer cell lines. The top of the figure shows classified and grouped PMDs (three classes) based on the average PMD methylation levels in PHH and their corresponding overall levels in HepaRG and HepG2, respectively. Note the intermediate status of HepaRG e.g. with a higher similarity to PHH in cluster1 (most highly methylated), an intermediate status in cluster 2 and a higher similarity to HepG2 in cluster 3 (most highly demethylated). The bottom shows the PMD wide changes in heterochromatic marks across the clusters defined by DNA-methylation. The inverse correlation to DNA-methylation is most obvious for HepaRG (class 1 and 3).

Along with the progressive loss of DNA-methylation in these three subclasses we observe a distinct gain of heterochromatic marks. The loss of DNA-methylation is correlated with the gain of heterochromatic marks (Figure 3C). The effect is most obvious in the HepaRG cell line which shows an intermediate level of PMD-methylation. Moreover, H3K36me3 is positively correlated with methylation across the gene body in the three subclasses (Figure S8). We confirmed this observation by calculating the average methylation across ChromHMM segments of HepaRG (Figure S10). PMDs associated with stronger transcription are higher methylated, on average, and marked by lower levels of heterochromatic marks.

We conclude that in immortalized cells a progressive erosion of DNA-methylation mainly in PMDs is linked to a substantial gain of heterochromatic marks. This is likely to be accompanied by differences in chromatin compaction and regulation in the immortalized cells with a prolonged proliferation. The conversion of PMDs, and sometimes of FMRs found in cancer tissues, into large unmethylated blocks as seen for HepG2, indicate that epigenetic changes found in model cell lines should be interpreted with great care, as they may reflect the properties more of the cell’s proliferation history and less of the cancer state or cell specific origin.

### Distinct heterochromatic signatures of PMDs predict replication timing

It has been shown that during cell division, late-replicating regions can become gradually demethylated [19] and long PMDs show widespread H3K9me3 marks bordered by H3K27me3, whereas shorter PMDs are enriched by H3K27me3 only [6]. So far, these features have not been thoroughly investigated and analyzed in an integrated fashion. Using replication timing data from the ENCODE project [20, 21] we clustered HepG2 hypomethylated/PMDs regions (longer than 300kb), by the k-means algorithm, into three clusters (see Methods and Figure 4A). These clusters display distinct histone modification and DNA-methylation patterns (Figure 4B). The first cluster (dark blue), consisting of the shortest PMDs, coincides with the early/middle S phase and is enriched for the two repressive marks H3K27me3 and H3K9me3. The second cluster (light blue) concurs with the middle/late S phase domains and comprises longer PMDs which are less highly enriched for H3K27me3 compared to cluster1. In the third cluster (yellow), PMDs extend over very long regions making up roughly 50% of the total PMDs/hypomethylated regions (Figure S12 and S13). These PMDs are strongly enriched for H3K9me3, bordered by H3K27me3 and coincide with the S4/G2 phases. Our findings suggest that combinations of heterochromatic marks and DNA-methylation define early, middle and late replication domains accompanied by a decrease in gene transcriptional activities during cell division (Figure 4B). We evaluated this result by predicting the three clusters using the three broad histone modification signals in PMDs, obtaining a high average prediction accuracy of 0.77 for HepG2 (0.81 for IMR90) (notice that this is a three-class prediction, details in Methods). The very late replicating regions in cluster 3 have the highest prediction accuracy, suggesting a distinct chromatin signature in the very late S4/G2 phase. These findings extend previous results [19] showing that demethylation in PMDs is accompanied by differences in chromatin signatures during the S-G2 phases.

**Figure 4.**
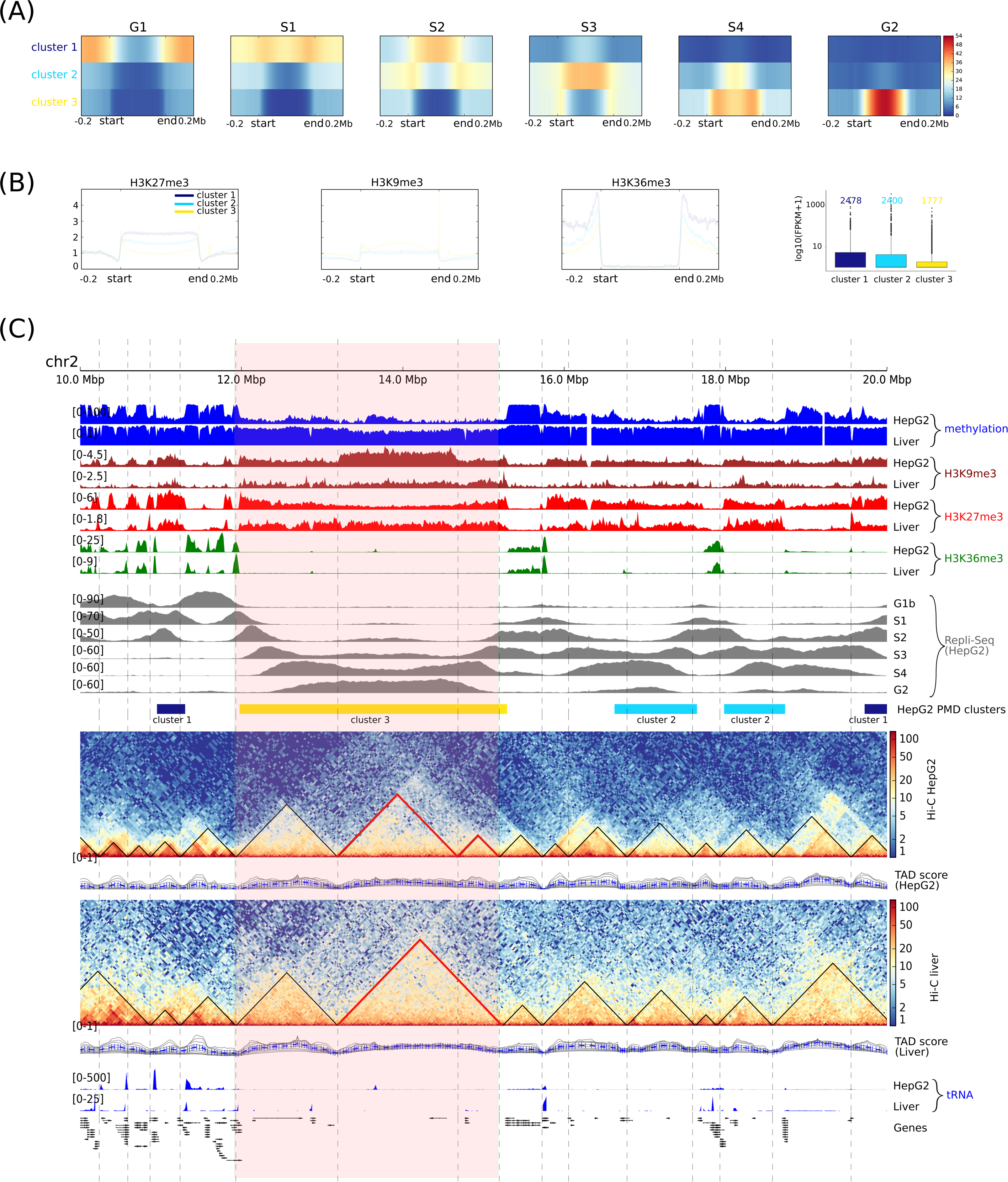
Distinct heterochromatin signatures of PMDs predict replication timing. (A) HepG2 PMDs are classified into three classes according to replication timing signals; cluster1 represents the early/mid S phase associated with PMDs-boundaries, cluster2 represents the middle/late S phase (S3/S4) and cluster3 represents the very late S phase (S4) and G2. (B) Epigenetic mark signatures across clusters in (A); H3K27me3 is highly enriched in cluster1 and in the PMD boundaries of cluster3 and less so in cluster2. H3K9me3 enrichment is similar in cluster1 and cluster2 and become more prominent in cluster3. The elongation mark H3K36me3 is depleted in all clusters. PMDs in cluster3 are the most demethylated and encompass the transcriptionally inactive genes. (C) Different epigenomic data tracks from chr2 shown in the following order: methylation profiles, H3K9me3, H3K27me3 and H3K36me3 histone marks of HepG2 and PHH, replication timing signals (G1-G2) of HepG2, clustered HepG2-hypomethylated/PMDs according to (A), Hi-C contact matrices of HepG2 and liver with the corresponding called TADs, tRNA and RefSeq genes. The highlighted region shows one long PMD, roughly 3Mb, extends over three TADs which are splitting according to H3K9me3 signal enrichment. Two of these TADs, marked in red, fuse into one TAD in the liver sample in agreement with the H3K9me3 signature.

To link these observations to high-order chromatin organization we generated Hi-C data for HepG2 and used available liver Hi-C data [22]. With the help of the HiCExplorer tool [23] we identified 3217 and 4021 TADs in liver and HepG2, respectively. We first observe that TADs of HepG2 are shorter than those in liver, on average. This is in agreement with Taberlay et.al. [24], who showed that cancer cells retain the ability to segment the genomes into generally smaller TADs due to establishment of new boundaries. As an example, Figure 4C (highlighted region) shows three TADs of HepG2, two of them fused into one TAD in the liver sample. An important observation is that the newly established TAD boundaries in HepG2 coincide with H3K9me3 boundaries but not with those of H3K27me3.

Our data show that the combined features of DNA-methylation and (hetero)chromatin modifications correlate with higher order chromatin structure and that they are determinants for phased replication. In the immortal cell line HepG2, marks such as H3K9me3 are more pronounced than H3K27me3, which we found to be more prominent in PHH (Figure S8, state 17_ReprPCWk). PMD are reflected in TAD chromatin domain structures indicating that PMDs are potential predictors of higher order chromatin structures.

## Discussion

This first comprehensive integrated analysis of primary human cells adds valuable novel insights into the structure and function of PMDs in human and mouse epigenomes. Building on our first report of PMD changes in primary T-cells [13], we extend the integrated approach across publicly available datasets to systematically analyze the features of PMDs in primary cells, primary tissues and immortalized cells from human and mouse. For our analysis we apply a new integrative “ChromH3M” approach which combines existing tools and represents an easy and straightforward method for analyzing and integrating a large cohort of WGBS data sets. This allowed us to define and compare PMDs across hundreds of WGBS samples, revealing a couple of intriguing new DNA-methylation properties with respect to genome organization, timing of DNA-replication and cell - type specific gene regulation.

First we find that PMDs comprise up to 75% of all epigenomes. However, only roughly 26% of the genome consists of PMDs that are conserved across all investigated cells. PMDs serve as excellent cell-type classifiers, and cells with functional similarities show a more similar PMD arrangement and topology, arguing for a shared developmental origin of lineage specific PMDs. We also observe that PMD methylation strongly changes in cultivated immortal cells.

In reminiscence to our previous interpretation of changes in proliferating B- and T-cells [13], we assume that the strong PMD changes in these immortal cells are also due to prolonged proliferation and cultivation.

Analyzing the PMD topology in more detail, we observe that the epigenomes of cells are partitioned into long regions of PMDs interspersed with short FMRs. These two classes of epi-domains show contrasting chromatin signatures. While PMDs are more heterochromatic and gene-poor regions, FMRs show strong transcriptional activity and enrichment of genes. This finding generalizes previous isolated observations reported by [4, 8, 10, 13, 25] to a number of different cancer types and cell lines. We also find cell-type specific changes from PMD to FMR, and vice versa, occurring in genomic regions that contain genes functionally enriched for cell type specific properties. This finding points towards a developmental control of PMD and FMR formation. The complete understanding which partitioning of FMRs and PMDs defines a precursor ground state of a cell type needs more investigation. Such knowledge will help understanding the role of epigenetic domains in cell differentiation.

We find that long PMDs have a lower density of protein coding genes, lincRNA and pseudogenes relative to the shorter PMDs (Figure S14). In general, protein coding genes are less highly expressed in long PMDs than in shorter PMDs and FMRs (Figure S14). We hypothesize that the long PMDs consists of more constitutive heterochromatin while the facultative heterochromatin can be found as shorter PMDs [26]. Shorter PMDs retain more epigenomic plasticity with more pronounced cell-type specific features.

PMDs can also be divided into different subclasses which are observed in different stages of DNA-replication. A hallmark of the late stages of replication (S4 and G2) is their length and the presence of the constitutive heterochromatic mark H3K9me3 together with a characteristic enrichment of H3K27me3 at the boundaries. On the other hand, the early/middle S phase (S1-3) PMDs are shorter and exhibit a higher overall proportion of H3K27me3. The length-dependent histone modification pattern is in agreement with previous findings in medulloblastoma [6]. The differences are strong enough to be used as predictors of the replication phases. Our results are consistent with a previous report [19] and extend its results by providing a detailed characterization of chromatin state and DNA-methylation at PMDs in relation to cell cycle. Moreover, the long PMDs associated with the late S4-G2 phases overlap with 56% of the bases within conserved PMDs. The shorter PMDs retain a greater variability, confirming our hypothesis that shorter PMDs possess epigenetically more less rigid heterochromatic structures than longer ones. This characteristic could be relevant for differentiation, cell-fate determination and or cell maturation processes.

To deepen our understanding about the cell type specific evolution of PMD (and FMR) structures, we compared the methylation landscape of human primary hepatocytes (PHH) with liver cancer tissue and hepatocellular carcinoma cell lines (HepaRG and HepG2). Overall, the methylome of primary liver cancer retained a PMD structure that is highly similar to primary cells, although with an overall mildly reduced level of methylation. In contrast, in cancer cell lines we observe a striking decrease of DNA methylation in such “mild” PHH_PMDs_, but not limited to these. The demethylated regions still retain the typical PMD histone marks although they are sometimes completely unmethylated. When such regions are considered as PMDs, the general PMD structure of cell lines is hepatocyte-like. The strong erosion of DNA-methylation is an example of how extensive cultivation leads to an enhancement of hepatocyte-like PMD demethylation. Such effects should be taken into account when drawing biological conclusions based on comparative epigenetic studies of *in vitro* cancer cell models. One cause of global demethylation in naive ESC is impaired DNA-methylation through the loss of H3K9me2, in combination with UHRF1 targeting replication loci [27]. Performing whole-genome hairpin sequencing will help to identify the hemimethylated sites that DNMT1 targets and to explore whether they harbor PMDs.

One way to consider global demethylation effects for the detection of differentially methylated region (DMR) was suggested in our recent study [13]. Here we screen for DMRs based on their deviation from the global methylation change rather than applying a fixed cutoff (for more details see [13]). DMRs were stratified over PMDs and FMRs such that many DMRs simply following the global demethylation could be excluded. In B-cells this procedure reduced the number of DMRs within PMDs tremendously (from 28014 to 8338 using adaptive filtering when comparing naive B-cells with plasma cells). On the other hand, DMRs, within PMDs, that gain methylation upon differentiation are increased (2811 DMRs in comparison to 95 retrieved by basic thresholding method). Genomic region enrichment analysis for such PMDs using GREAT [28] provides cell differentiation and development relevant terms (Figure S16). These findings demonstrate the advantage of stratifying DMRs according to increasing and decreasing of methylation in FMRs and PMDs, affording more insight into the biological role of the genes associated with these DMRs. Another way of using PMDs and FMRs is as a proxy for TADs and limiting the search space when annotating DMRs (or any regulatory regions) to genes.

A very recent study by Nothjunge, Stephan, et al. [29] showed that the establishment of B compartments precedes PMDs and define them during differentiation of cardiac myocytes. However, the underlying mechanisms are not yet clear. An interesting question would be whether establishment of B compartments also precedes the establishment of heterochromatic domains.

## Conclusion

We provide the first comprehensive analysis of PMDs for more than 190 human and mouse methylomes including more than 157 primary cell samples. Our analysis adds a new dimension to studying DNA methylation on a large-scale extending beyond the context of cis-regulatory elements that has been studied extensively. Our results show that PMDs are an excellent classifier of cellular origin and an indicator of the cellular proliferation history. In addition, PMDs heterochromatic histone mark signatures serve as an effective classifier for distinguishing early from middle and late replication domains. ChromH3M is an easy and straightforward framework for integrated analysis of large-scale WGBS data and can highlight specific combinatorial patterns of PMDs across large number of samples. PMDs are also a useful adjusting tool for detecting functional DMRs in highly proliferative cells. We believe that PMDs are a crucial epitopological signature beside their role in gene regulation. Our analysis reveals an important limitation in using cultivated cells for disease-associated epigenetic studies as they undergo strong changes in their epigenetic topology.

## Methods

### WGBS

Coverage and methylation fraction of human samples were downloaded from the Roadmap Epigenomic Project (http://egg2.wustl.edu/roadmap/data/byDataType/dnamethylation/WGBS). Blueprint data was downloaded from ftp://ftp.ebi.ac.uk/pub/databases/blueprint/data/homo-sapiens/GRCh38/ and then mapped to hg19 using liftOver from UCSC [30]. DEEP data was taken from previous studies [13, 31]. Bed files containing the coverage and methylation levels at CpG resolution from [29] were directly used in the analysis. We list all samples with the relevant sources in the additional file 1.

### MethylSeekR segmentation

All samples were segmented into partially methylated domain (PMDs), lowly methylated regions (LMRs) and unmethylated regions (UMRs) using the MethylSeekR tool [3]. The rest of the genome, excluding gaps as annotated by UCSC [30], was denoted as fully methylated regions (FMRs). We ran the tool with a coverage cutoff at five reads per CpG, methylation level threshold at 0.5 and maximum FDR of 0.05 for detection of hypomethylated regions, resulting in a threshold of at least four CpGs per LMR. The methylation levels of both strands were aggregated and weighted average methylation levels were plotted as box plots across PMDs.

### ChromH3M

In order to explore PMDs and find combinatorial patterns across samples, we binned the genome into 1kb windows and annotated each of them with 1 if the bin overlaps with a PMD and 0 otherwise across all samples. We used ChromHMM [15] to train this binarized signal with a 15-state HMM. We termed this method “ChromH3M”. The emission probabilities and states were hierarchically clustered using Euclidean distance and ward.D2 as an agglomeration method in the R environment [32]. The very same analysis was performed for LMRs and UMRs, respectively. To assess the uncertainty in the hierarchical clustering, we calculated an unbiased p-value (AU p-value) via multiscale bootstrap resampling (n=10000).

The normalized mean coverage of three broad histone marks (H3K27me3, H3K36me3 and H3K9me3), generated by the DEEP pipeline http://doi.org/10.17617/1.2W [33], were plotted genome-wide across the PMDs with proper flanking regions using deeptools [34]. The number of protein coding genes falling within PMDs was calculated, demanding a minimum of 80% of the gene length to be overlapping with the segment. A pseudocount of 1 was added to FPKM to avoid zeros in the box-plots.

The heatmap in Figure 3B was generated by binning the genome into 1kb windows and averaging the methylation levels across all samples resulting in roughly 280,000 windows which then were clustered by k-means into six clusters and annotated with methylSeekR segments. Samples were hierarchically clustered with ward.D2 and Euclidean distance. Sex chromosomes were excluded from the aforementioned analyses.

### Clustering of PHH PMDs and cancer cell lines

PHH PMDs shorter than 20kb were filtered and a matrix of methylation levels in 1kb windows across PHH, HepaRG and HepG2 was calculated after normalizing all PMDs to the same length of 150kb using deeptools [34]. The windows were clustered with k-means method into three clusters. H3K27me3, H3K9me3 and DNA-methylation signals were plotted along PMDs of each cluster using deeptools [34].

### Analysis of replication domains

Replication timing signals were downloaded from ENCODE project and used directly (detailes about this data are available from https://www.encodeproject.org/documents/50ccff70-1305-4312-8b09-0311f7681881/@@download/attachment/wgEncodeUwRepliSeq.html.pdf). A 2-state HMM was used to segment the HepG2 methylation profile into fully methylated and PMDs/hypomethylated regions using the “HiddenMarkov” R package [35], assuming that each

CpG may have one of the two states: foreground state with high methylation level and background state with low methylation level. Regions shorter than 300kb were filtered. The mean coverage of replication signals (G1, S1-S4 and G2) was calculated in 1kb bins across normalized (to 500kb length) and flanked PMDs (250kb up and down-stream) using deeptools [34]. PMDs were then clustered using k-means into three classes; early/middle S phase, middle/late S phase and late S/G2 phase. The mean coverage signals of H3K27me3, H3K9me3 and H3K36me3 and DNA-methylation levels were plotted across the PMDs of each class using deeptools [34]. The number of protein coding genes falling into each class was calculated, demanding 80% of the gene length to be within the PMD. FPKM values were plotted as box plots in log scale with pseudocount of 1 to avoid zeros.

For the prediction of replication domains, we built a multiclass classification model using the counts of reads of each histone mark in 1kb bins as predictors and the three aforementioned clusters as response at each PMD. We split the data into 75% training set and 25% test set. We trained the model with a random forest classifier and selected the model using 10-fold CV repeated five times. The prediction accuracy was calculated based on the confusion matrix between the predicted and the reference values. One-versus-all accuracy was calculated and then the average accuracy was calculated. This analysis was performed using the caret package https://github.com/topepo/caret/ in the R environment. The analysis of genomic regions regarding DMRs, with and without adjusting for the global DNA-demethylation was carried out using the GREAT tool [28]. GO analysis was done using DAVID [16, 17].

### Chromatin state segmentation

All Chip-Seq samples, listed in additional file1, were preprocessed starting from raw BAM files as follows: duplicate reads were removed using samtools version 1.3 with the filter "-F 1024". Regions of known artifacts ("blacklist regions") taken from the ENCODE project https://www.encodeproject.org/ [20], which we adapted to account for differences between ENCODE's hg19 and DEEP's hs37d5 assembly, were filtered out using bedtools version 2.20.1 with the subcommand "pairtobed" and the option "-type neither". After preprocessing, the filtered BAM files for all six histone marks plus Input were used as input for the chromatin state segmentation using ChromHMM version 1.11 (Java 1.7) with default parameters. We did not train a dedicated ChromHMM model for our dataset, but used the available ROADMAP 18-state model [36] to benefit from its biologically meaningful state labeling, which enabled us to immediately interpret the chromatin state maps in the context of this work.

### HepG2 Hi-C

HepG2 cells have been fixed for 10 minutes using 1% formaldehyde in D-MEM and quenched for 5 minutes in 125 mM glycine. After two PBS washes cells have been collected by scraping them off the plate and snap-frozen in liquid nitrogen. Hi-C experiments have been conducted as previously described [23], with the following modifications. Nuclei from cell pellets containing about 4 million of cells have been extracted by sonication [37] using the following parameters: 75W peak power, 2% duty factor, 200 cycles/burst, 180 seconds, using Covaris milliTubes and Covaris E220 sonicator. After nuclei permeabilization, chromatin has been digested overnight at 37 °C using HindIII high fidelity (80 units per million cells; R3104S, NEB). Biotin incorporation has been carried out at 37 °C for one hour in 300 μl volume using these reaction conditions: 50 mM of each nucleotide (dATP, dTTP, dGTP, biotin-14-dCTP, from Life Technologies, 19518-018), 8 U of Klenow (NEB, M0210L). Ligase mix has been added to each sample followed by 4 hours of incubation at room temperature under rotation. After nuclei lysis, protein digestion and overnight de-crosslink, DNA has been precipitated and sonicated to 100-600 bp. Biotinylated DNA has been pulled down as previously described. 100 ng of DNA bound to beads have been used for library preparation using a modification of the NEBNext Ultra DNA library preparation workflow (NEB, E7370). DNA bound to beads has been end-repaired, A-tailed, adaptor-ligated and USER-treated following manufacturer’s instruction. After a bead wash, DNA has been eluted from the beads by incubating at 98 °C for 10 minutes. Adaptor-ligated DNA has been PCR amplified using 7 PCR cycles. Libraries have been sequenced paired-end, with a read length of 75 bp, on the Illumina NextSeq 500 instrument.

### Hi-C data processing

Reads were mapped to the human reference genome hg19 (37d5) using bowtie2 [38], and then samtools [39] was used to convert the reads to BAM format. A matrix of read counts over the bins in the genome, considering the sites around the restriction site AAGCTT was built using the hicBuildMatrix function from HiCExplorer [23]. Ten bins were merged with hicMergeMatrixBins and then the matrix was corrected for GC bias and very low/high contact regions. To compute the TADs we first calculated the TAD scores by “hicFindTADs TAD_score” command with the following parameters “–minDepth 300000 –maxDepth 2000000 –step 70000” and then TADs were identified by “hicFindTADs find_TADs” command. The interaction matrix and other signal tracks were also visualized using HiCExplorer.

## List of abbreviations

PMDs: Partially methylated domains
FMRs: Fully methylated regions
LMRs: Lowly methylated regions
UMRs: Unmethylated regions
HMM: Hidden Markov Model
PHH: Primary human hepatocytes
TADs: Topological associated domains

## Declaration

### Ethics approval and consent to participate

Ethical approvals for human and mouse samples were present following standards outlined in DEEP, Blueprint and Roadmap. In our study we adhere to the rules defined by the data access committees.

### Consent for publication

Not applicable

### Data availability

WGBS data with the relevant sources used in this study are listed in additional file 1. WGBS mouse data (coverage and methylation calls) were downloaded from MethBase [40]. Chip-Seq and RNA-Seq DEEP data are available on the IHEC portal http://epigenomesportal.ca/ihec/grid.html. Liver Hi-C data was taken from [22] under the GEO accession number GSM1419086. HepG2 Hi-C data is available upon request. Replication timing signals of HepG2 and IMR90 were downloaded from the ENCODE project https://www.encodeproject.org/. All DEEP samples were uniformly processed as described in [13] according to DEEP pipelines (DOI: 10.17617/1.2W)

### Competing interests

Not applicable

### Funding

This work was mainly supported by the German Epigenome Programme (DEEP) of the German Federal Ministry of Education and Research (BMBF) (01KU1216F). JW and KN received partial support by the European Union’s Seventh Framework Programme (FP7/2007-2013) under grant agreement no 282510 – BLUEPRINT.

### Author’s contributions

study design (AS, KN, JW), integrative analysis (AS), chromatin state segmentation analysis (PE), HepG2 Hi-C data generation (LA, TM), Hi-C data analysis (AS, FR), HepaRG and HepG2 Chip-Seq data generation (KK), preparing Blueprint WGBS data in hg19 assembly (FM), providing hepatocyte samples (CC). AS, KN and JW wrote the manuscript with contributions from other authors. All authors read and accepted the final version of the manuscript.

## Acknowledgements

We are grateful to all colleagues from the IHEC consortium for making their data available. A full list of the investigators who contributed to the generation of the epigenomic data used in our study can be found under the respective homepages of the “NIH Epigenomics Roadmap”, “ENCODE”, the German epigenome consortium DEEP and BLUEPRINT (www.blueprint-epigenome.eu). We would like to thank Jasmin Kirch for technical support in NGS experiments.

